# Thymidine Phosphorylase Drives SARS-CoV-2 Spike Protein-Induced Lung Tumorigenesis

**DOI:** 10.64898/2025.12.14.694192

**Authors:** Cayleigh Wallace, Alex Gileles-Hillel, Amelia Cox, David Gozal, Wei Li, Hong Yue

**Author notes:** These two authors contribute equally (Cayleigh Wallace, Alex Gileles-Hillel, Wei Li, Hong Yue). **Corresponding Author:** Hong Yue, MD, PhD, Department of Biomedical Sciences, Joan C. Edwards School of Medicine at Marshall University, 1700 Third Avenue, Huntington, WV 25701, Or Wei Li, MD, PhD, FAHA, Department of Biomedical Sciences, Joan C. Edwards School of Medicine at Marshall University, 1700 Third Avenue, Huntington, WV 25701. **Author’s contributions:** Conception and design: HY, WL, DG. Data acquisition and analysis: CW, AGH, AC, WL, HY. Data interpretation: AGH, DG, WL, HY. Manuscript drafting: HY, WL. Critical revision of the manuscript: DG, AGH. Final approval of the version to be published: HY, WL. **Impact of this study:** Our findings provide mechanistic evidence that the SARS-CoV-2 Spike protein drives a profibrotic, pro-tumor microenvironment through TYMP-dependent signaling. This work offers a pathophysiological explanation for the emerging association between COVID-19 and increased lung cancer risk and identifies TYMP as a potential target for future preventive or therapeutic interventions.

## Abstract

**Rationale and Objectives:** COVID-19 survivors exhibit increased interstitial lung fibrosis, a known risk factor for lung cancer. We investigated whether SARS-CoV-2 Spike protein (SP)–induced lung injury and elevated thymidine phosphorylase (TYMP) promote lung tumorigenesis.

**Methods:** A TriNetX retrospective cohort analysis was combined with mechanistic studies in K18-hACE2^TG^ and K18-hACE2^TG^/*Tymp^−/−^* mice. Mice received intratracheal SP or control lysate followed by a urethane-induced lung cancer protocol. Lung injury, inflammation, thrombosis, fibrosis, STAT3 activation, cytokine profiles, and tumor burden were assessed. In vitro assays evaluated SP- and RBD-induced ACE2 processing.

**Results:** Propensity score–matched TriNetX cohorts demonstrated an increased lung cancer risk after COVID-19, particularly among current smokers (n = 166,807; RR 1.22; HR 1.50; P < .001). In mice, SP induced acute lung injury, neutrophil infiltration, and microthrombi, which were reduced in TYMP-deficient mice. SP markedly increased lung tumor incidence and aggressiveness, whereas TYMP deficiency reduced tumor formation from 50% to 18% of lung lobes. SP-induced STAT3 upregulation and collagen deposition were significantly attenuated in K18-hACE2^TG^/*Tymp^−/−^* mice. Cytokine profiling revealed a tumor-promoting, myeloid-dominant inflammatory milieu in K18-hACE2^TG^ mice, in contrast to a T cell–inflamed, anti-tumor profile in K18-hACE2^TG^/*Tymp^−/−^* mice. SP and RBD altered ACE2 processing, generating lower–molecular-weight fragments consistent with enhanced turnover.

**Conclusions:** SARS-CoV-2 SP drives lung injury, fibrosis, and tumorigenesis through a TYMP-dependent mechanism involving STAT3 signaling and inflammatory microenvironment remodeling. COVID-19 significantly increases lung cancer risk, especially in current smokers.

TYMP represents a potential therapeutic target to mitigate long-term pulmonary consequences of COVID-19.

## Introduction

COVID-19, caused by Severe Acute Respiratory Syndrome Coronavirus 2 (SARS-CoV-2) (1), has been diagnosed in over 778 million cases and led to 7.1 million deaths worldwide as of September 2025. Beyond the acute phase, survivors face substantial long-term complications, including interstitial pulmonary fibrosis (2), which occurs in approximately 25% of patients at three months and 14% at one year post-infection (3, 4). Because lung fibrosis is a well-established risk factor for lung cancer (5, 6), COVID-19 survivors may represent a population at heightened oncologic risk. Although COVID-19 vaccination has not been associated with increased lung cancer incidence (7, 8), recent studies report that SARS-CoV-2 infection can awaken dormant lung cancer cells and accelerate metastatic outgrowth (9). However, the molecular drivers linking SARS-CoV-2–related lung injury to cancer development remain largely unknown.

Emerging evidence suggests that the SARS-CoV-2 Spike Protein (SP) is intrinsically pathogenic. We and others have shown that SP is pro-inflammatory, pro-thrombotic, and capable of inducing a pro-fibrotic state in the lung (10, 11). Circulating SP has been detected in both infection and vaccination settings (12, 13), underscoring the need to determine whether SP directly drives lung fibrotic remodeling and contributes to oncogenesis.

Thymidine phosphorylase (TYMP) is a cytoplasmic and nuclear protein with pro-angiogenic and signaling functions (14–16). TYMP expression is elevated in numerous pathological conditions, including atherosclerosis, cancer, and diabetes mellitus, and higher TYMP levels are associated with poor cancer prognosis (14, 17). TYMP is significantly increased in the plasma, neutrophils, macrophages, and lungs of COVID-19 patients (18–20), correlating with thrombotic events, inflammation, and respiratory symptoms. TYMP also participates in fibrotic processes in liver cancer and urinary tract disease (21, 22), and elevated TYMP expression in non-cancerous tissues predicts multifocal hepatocellular carcinoma (23). Together, these observations suggest that SARS-CoV-2-induced TYMP elevation may promote post-COVID lung fibrosis and increase lung cancer susceptibility.

In this study, using a TriNetX Research Network human cohort and complementary mouse models, we tested this hypothesis and demonstrate that TYMP promotes SARS-CoV-2 SP–enhanced lung cancer development.

## Materials and Methods

### Human study to examine the association between SARS-CoV-2 infection and cancer development

We conducted a retrospective cohort study using deidentified electronic health records from the TriNetX Research Network. Because only aggregated, anonymized data were assessed, this study did not meet the definition of human subjects research under U.S. federal regulations and did not require Institutional Review Board approval. Patients were stratified by smoking status (current, former, never) and categorized by documented SARS-CoV-2 infection and vaccination status. Patients with recorded outcomes before the prespecified observation window were excluded.

Two sets of comparisons were performed within each smoking stratum.

1. Primary exposure analysis (COVID-19 vs no COVID-19):

- Current smokers: COVID-19 (largely unvaccinated) vs no COVID-19 (vaccinated)
- Former smokers: COVID-19 (unvaccinated) vs no COVID-19 (vaccinated)
- Never smokers: COVID-19 (vaccinated) vs no COVID-19 (vaccinated).
2. Effect-modification analysis (vaccination vs unvaccinated among COVID-19-positive patients):

- Current smokers, former smokers, and never smokers analyzed separately.

TriNetX implemented 1:1 propensity score matching (PSM) for each comparison on age, sex, race, and ethnicity. Covariate balance was assessed using standardized differences, and outcome were evaluated in the matched cohorts. The primary outcome was incident lung cancer (ICD-10-CM C34). Secondary outcomes included incident oral cancer and bladder cancer (ICD-10-CM C67).

We calculated absolute risks, risk differences (RDs), risk ratios (RRs), and odds ratios (ORs).

Time-to-event analyses employed Kaplan–Meier curves, log-rank tests, and Cox proportional hazards models with proportionality checks. Two-sided P values <0.05 were considered statistically significant. All analyses were performed within the TriNetX platform, and primary estimates were derived from matched populations with verified covariate balance.

### Generation of SARS-CoV-2 Spike Protein

Crude SARS-CoV-2 SP–containing cell lysate (SP) and control cell lysate transfected with the empty plasmid pcDNA3.1 (P3.1) were prepared as previously described (10). Lysates were bulk-generated, aliquoted into single-use tubes containing 500 µg total protein in 50 µL PBS and immediately stored at -80°C.

### Mouse model of lung cancer development

K18-hACE2^TG^ mice (Strain: 034860, The Jackson Laboratory) and K18-hACE2^TG^/*Tymp^-/-^* mice generated in our recent study (10) were used. Mice were anesthetized with ketamine/xylazine (100/10 mg/kg) and intubated with an 18-gauge needle catheter. A total of 500 µg SP or P3.1 cell lysate in 50 µL PBS was administered intratracheally over one minute, after which mice were allowed to recover from anesthesia.

Mice were randomly allocated into two groups. In group one, whole blood was collected from the inferior vena cava 24 hours later using 0.109 M sodium citrate as an anticoagulant, and whole blood cell counts were obtained using a Hemavet 950SF analyzer. Mice then euthanized, lungs were inflated with 30 cm H2O pressure, and tracheas were ligated. Lungs were fixed in 10% formalin, paraffin-embedded, and sectioned for histological examination.

In group two, mice received intraperitoneal urethane injections at 1 g/kg starting the next day after SP or P3.1 administration, once weekly for eight weeks, followed by a 20-week latency period (24, 25). Whole blood was collected from the inferior vena cava, and plasma was isolated by centrifugation. Lungs were processed as in the 24-hour study, fixed in formalin for 48 hours, and lung tumors were examined visually and microscopically.

Mice had *ad libitum* access to standard laboratory rodent diet and water. Both males and females were used in this study. Ethical approval was obtained from the Institutional Animal Care and Use Committee (IACUC) of Marshall University (IACUC#: 1033528, PI: Wei Li). All procedures adhered to the NIH Guide for the Care and Use of Laboratory Animals.

### Determine how SARS-CoV-2 Spike Protein affects ACE2 expression

BEAS-2B cells were treated with P3.1 or SP for 24 hours. Cells were then lysed and processed for Western blotting analysis of ACE2 expression following standard protocols.

To further evaluate how SARS-CoV-2 SP affects ACE2 function, COS-7 cells were co-transfected with plasmids encoding human ACE2 and either C9-tagged SP or GFP-tagged S1 receptor binding domain (RBD). Cell lysates were collected for Western blot assay of ACE2, SP, and RGD expression, with pan-actin used as the loading control. All antibodies used in this project were listed in **Supplemental Table 1**

### Histology examination

Formalin-fixed and paraffin-embedded lung tissues were sectioned into 6 µm. Hematoxylin and eosin (H&E) and Masson’s Trichrome staining were performed to assess lung structure, lung tumors, and fibrosis. Standard single or double immunohistochemistry (IHC) was conducted using antibodies as detailed in the Results section.

### Mouse plasma cytokine array

Twenty microliters of plasma from each mouse were pooled from six randomly selected K18-hACE2^TG^ mice and six K18-hACE2^TG^/*Tymp^-/-^* mice that received SARS-CoV-2 SP. The pooled plasma was then used to measure cytokine levels using the Proteome Profiler Mouse Cytokine Array Kit, Panel A (Catalog # ARY006, R&D Systems, Inc. Minneapolis, MN). Membrane images were scanned, and spot intensities were quantified using ImageJ. Data were presented as log2(fold change of K18-hACE2^TG^/K18-hACE2^TG^/*Tymp^-/-^*). A value of one was assigned to any spot that was undetectable in one strain but detectable in the other to allow for mathematical calculations.

### Statistics

Data were analyzed using GraphPad Prism (version 10.4.2) and were expressed as mean ± SEM. Statistical analyses were performed using a two-tailed Student’s t-test, Mann–Whitney test, or One or Two-way ANOVA, as appropriate. Uncorrected Fisher’s LSD test was used as the post hoc analysis for Two-way ANOVA. A p-value of ≤ 0.05 was considered statistically significant.

## Results

### 1. SARS-CoV-2 infection increases the incidence of lung cancer in humans

Using the retrospective cohort study through the TriNetX Research Network, we analyzed the correlation between COVID-19 and cancer development. **Supplemental Table 2** shows the size of the matched cohort and its demographics. We found that prior COVID-19 infection is significantly associated with an increased risk of incident lung cancer, with the highest hazard observed among current smokers, followed in decreasing order by former smokers and non-smokers (**Table 1**). In the same cohort, however, oral and bladder cancers showed no consistent increase in risk with prior COVID-19 (**Table 2**). These data support recent findings that SARS-COV-2 infection can awaken dormant lung cancer cells and promote tumor development (9). Clarifying the underlying mechanism is therefore urgently needed.

**Table 1.** Lung cancer risk and time-to-event results in matched cohorts.

| Smoking status & comparison | Risk A | Risk B | RD (A–B) | RR (95% CI) | HR (95% CI) | Log-rank P |
| --- | --- | --- | --- | --- | --- | --- |
| Current, COVID-19 vs No COVID-19 | 1.7% | 1.4% | +0.3% | 1.22 (1.15–1.29) | 1.50 (1.42–1.58) | <.001 |
| Former, COVID-19 vs No COVID-19 | 1.5% | 1.2% | +0.3% | 1.24 (1.20–1.29) | 1.60 (1.54–1.66) | <.001 |
| Never, COVID-19 vs No COVID-19 | 0.21% | 0.18% | +0.0% | 1.16 (1.10–1.21) | 1.42 (1.35–1.49) | <.001 |
| Current with COVID-19, Unvaccinated vs Vaccinated | 1.9% | 1.8% | +0.1% | 1.08 (1.03–1.13) | 1.02 (0.97–1.07) | .42 |
| Former with COVID-19, Unvaccinated vs Vaccinated | 1.5% | 1.3% | +0.2% | 1.19 (1.15–1.23) | 1.14 (1.10–1.17) | <.001 |

**Abbreviations:** A indicates the exposure cohort listed first in each comparison; B, the comparator. RD, risk difference; RR, risk ratio; HR, hazard ratio. Outcomes exclude individuals with the event before the analysis window. PSM indicates propensity score matching.

**Table 2.** Oral cancer (secondary outcome) in matched cohorts.

| Smoking status & comparison | Risk A | Risk B | RD (A–B) | RR (95% CI) | HR (95% CI) | Log-rank P |
| --- | --- | --- | --- | --- | --- | --- |
| Current, COVID-19 vs No COVID-19 | 0.27% | 0.29% | –0.0% | 0.92 (0.81–1.05) | 1.15 (1.01–1.31) | .03 |
| Former, COVID-19 vs No COVID-19 | 0.28% | 0.36% | –0.1% | 0.78 (0.72–0.84) | 1.01 (0.94–1.09) | .73 |
| Never, COVID-19 vs No COVID-19 | 0.08% | 0.08% | –0.0% | 0.97 (0.90–1.04) | 1.23 (1.14–1.33) | <.001 |
| Current with COVID-19, Unvaccinated vs Vaccinated | 0.28% | 0.27% | +0.0% | 1.04 (0.92–1.17) | — | .56 |
| Former with COVID-19, Unvaccinated vs Vaccinated | 0.27% | 0.24% | +0.0% | 1.13 (1.05–1.22) | 1.09 (1.01–1.18) | .03 |

**Abbreviations:** A indicates the exposure cohort listed first in each comparison; B, the comparator. RD, risk difference; RR, risk ratio; HR, hazard ratio. Outcomes exclude individuals with the event before the analysis window. PSM indicates propensity score matching.

### 2. Intratracheal delivery of SARS-CoV-2 SP induces lung injury, inflammation, and microthrombus formation

Given that COVID-19 is primarily a respiratory disease, we examined the effect of intratracheal SP administration on lung inflammation and thrombus formation in K18-hACE2^TG^ and K18-hACE2^TG^/*Tymp^-/-^* mice. As shown in **Supplemental Table 3**, intratracheal SP administration did not significantly affect whole blood cell counts at 24 hours.

The receptor for advanced glycated end products (RAGE), expressed by bronchiolar epithelial cells, type II alveolar cells, and alveolar macrophages (26), plays an important role in lung inflammation and is considered a marker of acute lung injury. As shown in **Fig. 1A**, inhalation of either P3.1- or SP-containing cell lysate induced significant lung injury; however, SP caused more severe injury, which was attenuated by TYMP deficiency.

**Figure 1.**
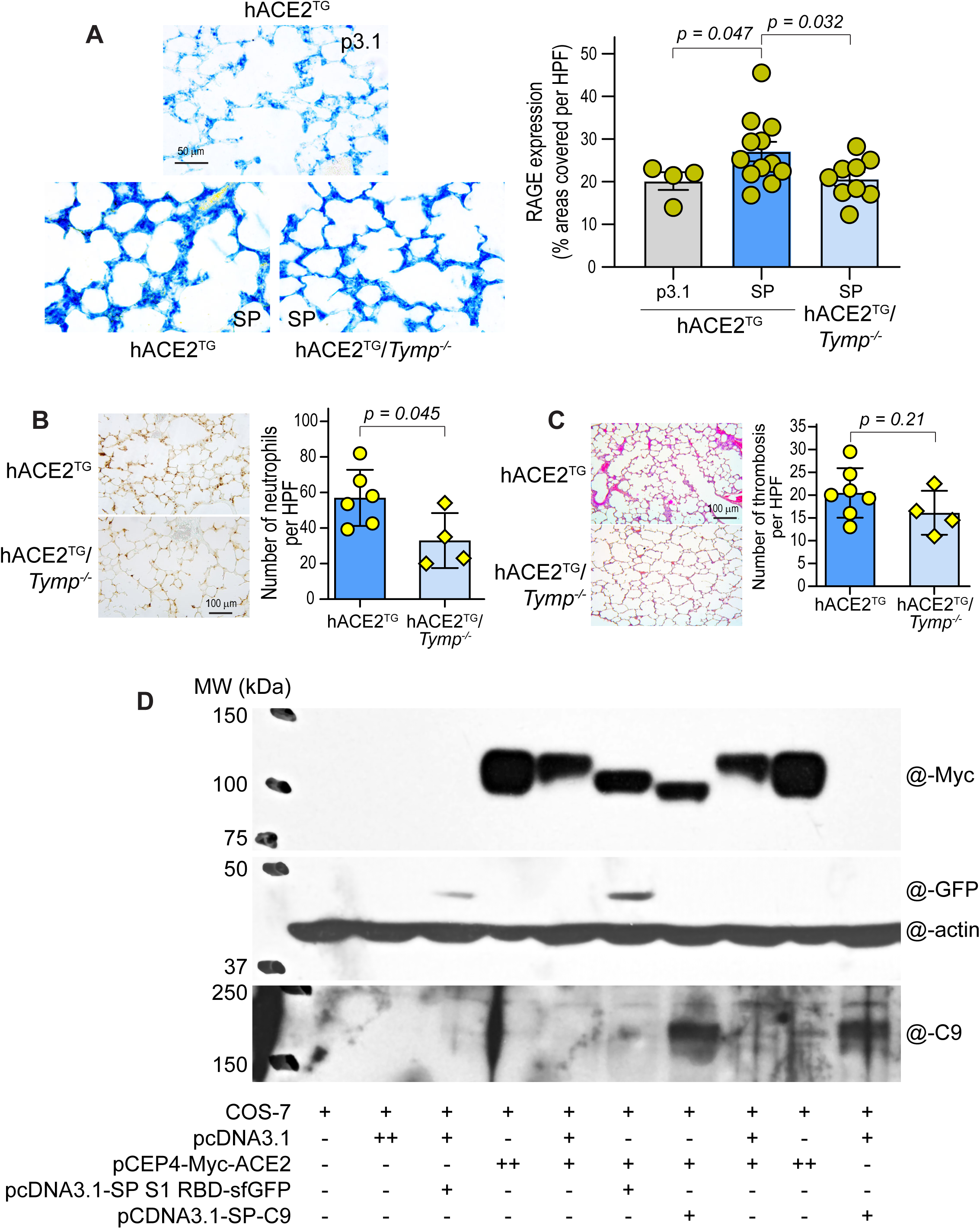
SARS-CoV-2 spike protein (SP) induces lung injury, inflammatory cell infiltration, microthrombus formation, and ACE2 shedding. K18-hACE2^TG^ (hACE2^TG^) and K18-hACE2^TG^/*Tymp^-/-^* (hACE2^TG^/*Tymp^-/-^*) mice received intratracheal administration of 500 µg COS-7 cell lysate containing either SP or control (P3.1). Lungs were harvested 24 hours later for histological analysis. A. Lung injury was assessed by immunohistochemical staining of the receptor for advance glycated end products (RAGE), visualized with Vector Blue. No counterstaining was performed. Images were analyzed using ImageJ, and data are presented as the percentage of area covered per high-power field (HPF, 40x) (right panel). B. Immunohistochemical staining for myeloperoxidase (MPO) was performed as a marker of neutrophil infiltration. MPO-positive cells (brown) were quantified using ImageJ. C. Hematoxylin & Eosin staining was used to evaluate lung morphology. Microthrombi were counted visually with the assistant of ImageJ. D. COS-7 cells were co-transfected with the indicated plasmids in various combinations using FuGENE® 6 Transfection Reagent. All plasmids were obtained from Addgene. Cells were harvested 40 hours after transfection and analyzed by Western blot using the indicated antibodies. Antibody dilutions are listed in Supplemental Table 1. Data represents two independent biological replicates.

Acute lung injury is accompanied by an influx of neutrophils into the interstitial and bronchioalveolar space (27). We therefore stained lung neutrophils using myeloperoxidase as a marker. Neutrophil infiltration was significantly reduced in K18-hACE2^TG^/*Tymp^-/-^* mice 24 hours after SP exposure (**Fig. 1B**), consistent with the reduced lung injury observed in these mice. In contrast, CD68-positive macrophage levels were similar between P3.1 and SP treatments and between genotypes (**Supplemental Figure 1**).

Microthrombi were detected in the lungs of all mice, including controls receiving P3.1 lysates (data not shown). However, SP inhalation induced larger and more extensive thrombus formation, which was attenuated in TYMP-deficient mice (**Fig. 1C**). These findings support our previous observations that SARS-CoV-2 SP enhances inflammation and thrombosis (10), contributing to lung injury.

Reportedly, binding of the SARS-CoV-2 SP or its RBD to ACE2 induces ACE2 internalization and degradation, resulting in reduced surface ACE2 expression (28, 29). Although BEAS-2B cells express a low level of ACE2, we did not detect ACE2 protein under our experimental conditions in cells treated with P3.1, SP, or vehicle (data not shown). We therefore co-transfected human ACE2 together with SP or RBD into COS-7 cells, which lack endogenous ACE2, to examine their effects on ACE2 protein processing.

Unexpectedly, instead of observing uniform ACE2 degradation, we found that co-expression of SP or RBD generated distinct lower-molecular weight Myc-tagged ACE2-immunoreactive bands (**Fig. 1D**), with SP producing a further smaller fragment. Because the Myc-tag is positioned at the ACE2 N-terminus, the presence of these smaller Myc-positive fragments suggests that SP and RBD induced ACE2 processing that removes portions of the C-terminal region, consistent with that ACE2 is a target of intramembrane proteolysis of γ-secretase, releasing a soluble ACE2 C-terminal fragment (30).

### 3. Intratracheal delivery of SARS-CoV-2 SP increases lung cancer incidence

Given that lung cancer incidence is significantly increased post-COVID-19 (9), we examined the long-term consequences of SARS-CoV-2 SP–induced lung injury by subjecting mice that received P3.1 or SP treatment to a urethane-induced lung cancer model (**Fig. 2A**). In K18-hACE2^TG^ mice that received SP, tumors (**Fig. 2B**) were observed in 10 of 20 lung lobes examined. In contrast, only one tumor was identified among the 10 lobes from mice treated with P3.1 control cell lysate (**Fig. 2C**; Fisher’s exact test-*P* value:0.0485), indicating that SP exposure significantly promoted lung tumor development. Whereas tumors were detected in 50% of the lobes examined in K18-hACE2^TG^ mice, only nine tumors were found among 50 lobes (18%) in K18-hACE2^TG^/*Tymp^-/-^* mice (**Fig. 2D**), indicating that TYMP deficiency markedly reduces SP-induced lung tumorigenesis. Histological analysis revealed that SP-enhanced tumors were more aggressive, as evidenced by their large tumor size (**Fig. 2E vs. 2F**). Unexpectedly, all tumors found in SP-treated mice were positive for p40 staining (**Fig. 2G vs. 2H**), a marker of lung squamous cell carcinoma (31, 32), which is not commonly observed in the urethane-induced lung cancer model that typically shows a bronchioloalveolar adenoma phenotype (33).

**Figure 2.**
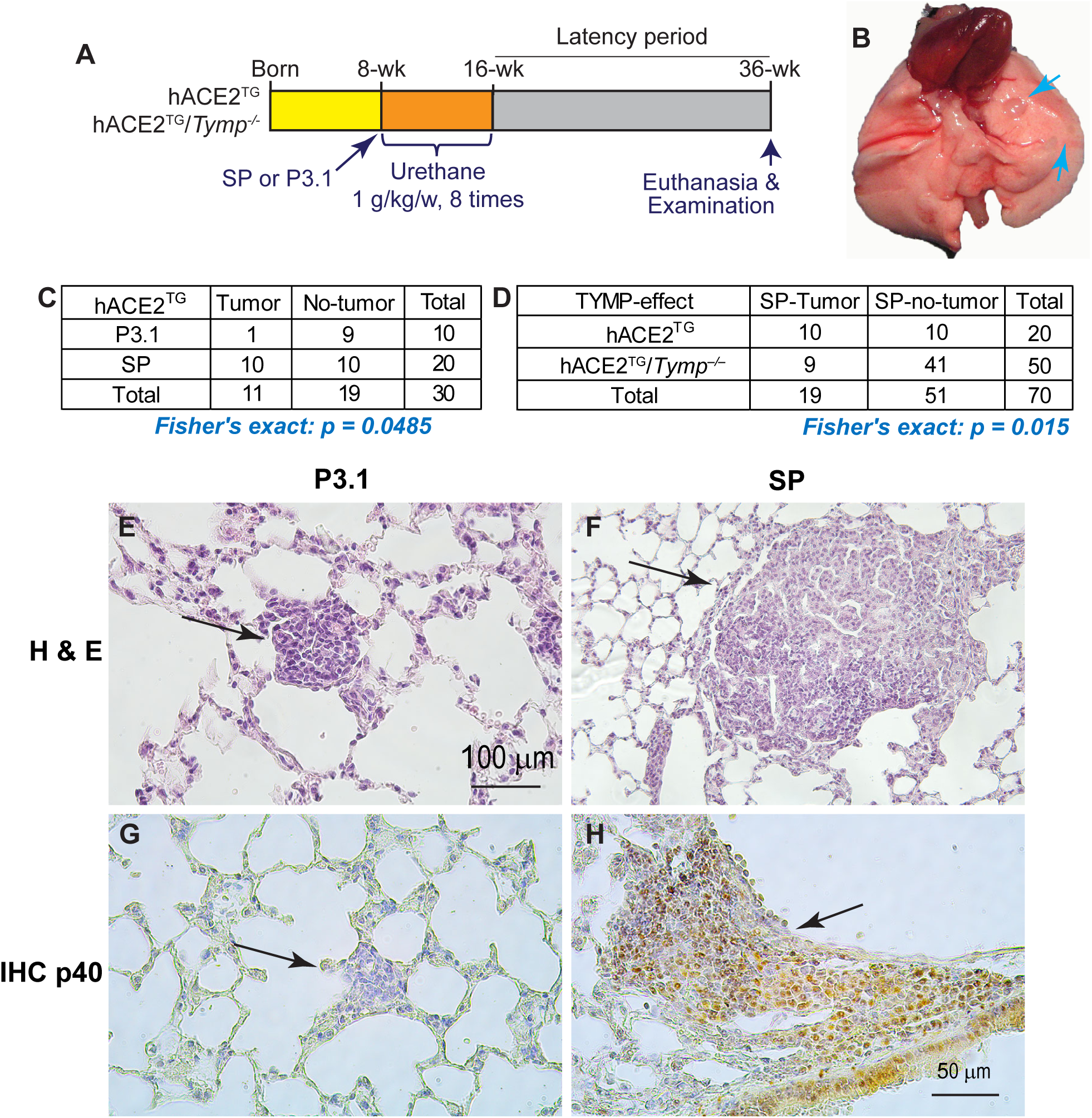
Intratracheal delivery of SARS-CoV-2 Spike protein (SP) increases lung cancer incidence. **A.** Schematic illustration of the urethane-induced lung cancer model. **B.** Representative gross images of lung cancer. **C**. SP-enhanced lung tumor incidence in K18-hACE2^TG^ (hACE2^TG^) mice receiving SP or P3.1 control lysate. **D.** Comparison of SP-enhanced lung tumor incidence between K18-hACE2^TG^ (hACE2^TG^) and K18-hACE2^TG^/*Tymp^-/-^*(hACE2^TG^/*Tymp^-/-^*) mice. **E & F**. Hematoxylin & Eosin staining with representative histological images of lung tumors. **G & H**. Immunohistochemical staining for p40, a marker of lung squamous cell carcinoma. Brown staining indicates p40-positive cells.

We previously reported that TYMP inhibits vascular smooth muscle cell (VSMC) proliferation (34, 35), whereas TYMP deficiency does not affect angiogenesis in mice (36). Using CD31 and α-smooth muscle actin (α-SMA) double immunohistochemistry, we further confirmed that TYMP deficiency also does not affect lung angiogenesis or arteriogenesis in K18-hACE2^TG^ mice (**Supplemental Figure 2**). These findings suggest that the differences in lung cancer incidence observed are not attributable to TYMP’s pro-angiogenic effect or its influence on VSMC function.

### 4. TYMP amplifies STAT3 signaling and drives a profibrotic, pro-tumor microenvironment

We recently demonstrated that both SARS-CoV-2 RBD and SP significantly increased STAT3 phosphorylation at Y705, which was markedly attenuated by siRNA-mediated knocking down of TYMP (10). This finding aligns with our previous observation that TYMP overexpression enhances STAT3 activation (37). Moreover, several studies suggest that STAT3 plays a potential role in COVID-19 pathophysiology (38). To determine if TYMP-mediated, SP-enhanced STAT3 activation contributes to cancer development, we stained lung tumor sections for Y705- phosphorylated STAT3 and total STAT3. As shown in **Fig. 3A**, while Y705-STAT3 was widely stained in lung tissues regardless of tumor presence, total STAT3 was predominantly localized within tumors.

**Figure 3.**
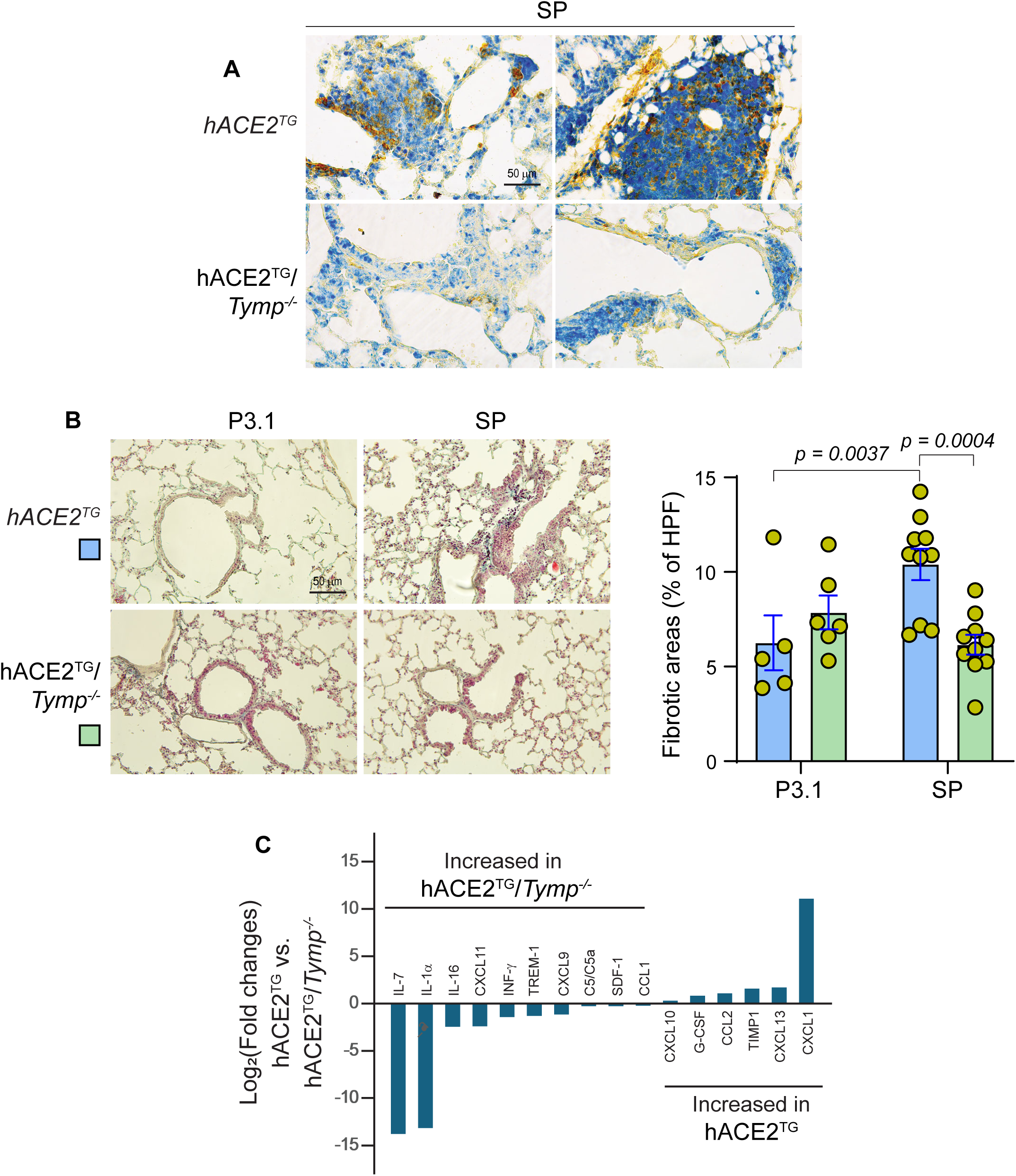
TYMP amplifies STAT3 signaling and drives a profibrotic, pro-tumor microenvironment. **A.** Lung sections from SP-treated K18-hACE2^TG^ (hACE2^TG^) and K18-hACE2^TG^/*Tymp^-/-^* (hACE2^TG^/*Tymp^-/-^*) mice were stained for Y705-phosphorylated STAT3 (Blue) and Total STAT3 (Brown). **B**. Trichrome staining was performed on lung sections from mice subjected to the urethane-induced lung cancer protocol. Images were analyzed using ImageJ, and fibrotic areas were quantified by measuring blue-stained regions obtained through color deconvolution. Quantitative data are presented as bar grafts (right panel). Statistical analyses were performed using Two-way ANOVA followed by Uncorrected Fisher’s LSD post hoc testing.

Given that STAT3 activation is associated with fibroblast-to-myofibroblast transition, collagen deposition, and fibrosis both in vitro and in vivo (39), we next evaluated lung fibrosis in mice subjected to the lung cancer model using Trichrome staining. A two-way ANOVA revealed a significant Genotype x Treatment interaction (F(1,27) = 10.67, *p* = 0.003), indicating that the effect of treatment differed by genotype. Neither Genotype (*p* = 0.185) nor Treatment (*p* = 0.150) showed a significant main effect. *Post hoc* comparisons demonstrated that, compared with P3.1-treated controls, SP markedly increased lung fibrosis in K18-hACE2^TG^ mice (**Fig. 3B**). In contrast, this SP-induced fibrosis was significantly attenuated in K18-hACE2^TG^/*Tymp^-/-^*mice (difference of differences = −5.848, 95% CI: −9.521 to −2.175). Together, these findings suggest that the interaction between SARS-CoV-2 SP and host cells establishes a microenvironment that is concurrently profibrotic and pro-carcinogenic.

We previously reported that TYMP deficiency alters the expression of several cytokines (40), which may influence immune disorders. To assess whether the interaction between SP and TYMP modulates cytokine expression, we pooled plasma from 6 K18-hACE2^TG^ and 6 K18-hACE2^TG^/*Tymp^-/-^* mice treated with SP and conducted a cytokine array analysis. As shown in **Fig. 3C and Supplemental Figure 3**, compared with K18-hACE2^TG^, K18-hACE2^TG^/*Tymp^-/-^*mice exhibited increased levels of Th1-associated mediators (IFN-γ, IL-7), CXCR3 ligands (CXCL9, CXCL11), and T-cell–attracting chemokines (IL-16, CCL1), consistent with a T-cell–inflamed, anti-tumor microenvironment. In contrast, K18-hACE2^TG^ mice showed elevated G-CSF, CCL2, CXCL1, TIMP-1, and CXCL13, characteristic of a myeloid-dominated, tumor-promoting inflammatory state, aligning with the higher incidence of lung tumors observed in K18-hACE2^TG^ mice.

## Discussion

In this study, we provide convergent human and experimental evidence that SARS-CoV-2 infection increases the risk of lung cancer and identify TYMP as a previously unrecognized molecular driver of Spike Protein (SP)–induced lung injury, fibrosis, and tumorigenesis. By integrating large-scale clinical data with mechanistic mouse models, we demonstrate that TYMP amplifies SP-triggered inflammation and fibrotic remodeling, facilitates STAT3 expression, and shapes a tumor-permissive immune microenvironment. These findings establish TYMP as a central pathogenic mediator linking SARS-CoV-2 respiratory injury to increased lung cancer risk.

Our analysis of a large, demographically matched cohort from the TriNetX Research Network revealed a significant increase in lung cancer incidence among individuals with prior COVID-19 infection. This effect was strongest among current smokers, followed by former smokers and non-smokers, consistent with the idea that preexisting epithelial injury or chronic inflammation sensitizes the lung to SARS-CoV-2–induced oncogenic stimuli (41, 42).

Importantly, this increased risk was not observed for oral or bladder cancers, indicating that the association may be organ-specific and related to mechanisms unique to the lung microenvironment. Interestingly, we previously found that TYMP was not expressed in several tongue cancer cell lines, including HSC3 and SCC-9 (14). These findings align with recent reports that SARS-CoV-2 may selectively awaken dormant lung cancer cells and promote rapid tumor progression (9), underscoring the need to identify biological processes that couple viral exposure to oncogenesis.

Because SARS-CoV-2 SP is the first viral protein to interact with host cells, we investigated whether SP alone is sufficient to recapitulate lung pathological features associated with (and potentially contributing to) tumor development. Intratracheal SP delivery caused marked acute lung injury, characterized by heightened RAGE staining, neutrophil infiltration, and extensive microthrombus formation. These effects were significantly attenuated in TYMP-deficient mice, indicating that TYMP amplifies SP-triggered inflammatory and thrombotic responses. Although macrophage infiltration did not differ between genotypes, the robust neutrophilic influx in K18-hACE2^TG^ mice highlights TYMP’s selective role in shaping acute inflammatory responses.

Neutrophils are known to promote DNA damage, secrete pro-tumorigenic proteases, and generate neutrophil extracellular traps, all of which may contribute to tumor initiation. Thus, the early inflammatory landscape induced by SP, and modulated by TYMP, could establish a foundation for subsequent malignant transformation.

Using a urethane-induced lung cancer model, we found that prior SP exposure significantly increased tumor incidence and aggressiveness. We used a urethane dose of 1 g/kg, which has been reported to induce lung tumors with a 60-80% incidence (24, 25). However, in K18-hACE2^TG^ mice that received P3.1, we observed only ∼10% tumor incidence. This discrepancy may be explained by the fact that ACE2 is known to inhibit proliferation, metastasis, invasion, and angiogenesis of various tumor types, including lung cancer (43, 44). SP- or RBD-associated ACE2 shedding or degradation may diminish this protective effect of ACE2.

Remarkably, tumors in SP-treated mice exhibited squamous cell carcinoma morphology and strong p40 positivity, a phenotype atypical for urethane-induced adenomas. These findings suggest that SP and TYMP interaction reprograms the epithelial injury–repair process toward squamous metaplasia and malignant transformation.

Strikingly, TYMP deficiency dramatically reduced tumor number and size, indicating that TYMP is essential for SP-enhanced lung tumorigenesis. Importantly, the reduction in tumor incidence was not attributable to impaired angiogenesis or defects in VSMC function, as TYMP-deficient mice showed normal pulmonary vascular architecture (Supplemental Figure 2). Instead, our findings point to TYMP’s regulatory role in inflammatory signaling and fibrotic remodeling as key determinants of tumor promotion.

STAT3 is a master regulator of inflammation, fibrosis, and cancer progression (39). Building on our prior observation that SP enhances STAT3 Y705 phosphorylation in BEAS-2B cells in a TYMP-dependent manner (10), we now show that phospho-STAT3 is broadly detected across lung tissues, whereas total STAT3 is primarily enriched within tumors from K18-hACE2^TG^ mice. STAT3 activation promotes fibroblast–myofibroblast transition, extracellular matrix deposition, and epithelial plasticity (45). Our previous study further demonstrated that unphosphorylated STAT3 also possesses pathophysiological functions (46); its expression is increased in TYMP-overexpressing VSMCs and suppresses their proliferation (37). Together, these findings suggest that TYMP-mediated STAT3 signaling serves as a mechanistic bridge linking SP-induced injury to fibrosis and carcinogenesis.

Consistent with this hypothesis, we found that SP markedly increased lung collagen deposition and fibrotic remodeling in K18-hACE2^TG^ mice, whereas fibrosis was significantly attenuated in K18-hACE2^TG^/*Tymp^-/-^* mice. Because pulmonary fibrosis is a well-recognized risk factor for lung cancer, these results support a tangible pathogenic framework, which creates a tumor-permissive niche.

Cytokine profiling revealed fundamentally different inflammatory landscapes between SP-treated K18-hACE2^TG^ mice and K18-hACE2^TG^/*Tymp^-/-^*mice. SP-exposed K18-hACE2^TG^ mice exhibited elevated G-CSF, CCL2, CXCL1, TIMP-1, and CXCL13, factors associated with myeloid recruitment, angiogenesis, extracellular matrix remodeling, and tumor promotion. In contrast, K18-hACE2^TG^/*Tymp^-/-^* mice showed enhanced Th1 cytokines (IFN-γ, IL-7), CXCR3 ligands (CXCL9, CXCL11), and T-cell–attracting chemokines (IL-16, CCL1) (47, 48), indicative of an inflamed, anti-tumor immune environment. Overexpression of certain forms of IL-1*α* has been shown to lead to tumor regression in some experimental models (49). Thus, TYMP not only amplifies SP-induced injury and fibrosis but also orchestrates an immunologic shift toward tumor-supportive inflammation. This dual role, structural and immunologic, may explain the profound reduction in tumor development observed in TYMP-deficient mice.

Taken together, our findings support a mechanistic model in which SARS-CoV-2 SP initiates lung injury, neutrophilic inflammation, and microthrombosis; induces sustained TYMP expression; and drives STAT3-mediated fibrosis and immune dysregulation. This creates a microenvironment conducive to malignant transformation or expansion of previously dormant tumor cells. Clinically, these results suggest that COVID-19 survivors may benefit from long-term lung cancer surveillance, particularly those who are current and former smokers. TYMP is a compelling therapeutic target. Tipiracil, an FDA-approved TYMP inhibitor known to inhibit thrombosis (50), may warrant investigation for preventing post-COVID fibrosis or mitigating cancer risk. STAT3-directed therapies may be beneficial in patients with persistent post-COVID lung injury.

Our study is limited by reliance on retrospective clinical data, which obviously precludes any inferences of causality. Prospective longitudinal studies are necessary to quantify cancer risk over time after infection. Mechanistically, the precise molecular pathways downstream of TYMP remain to be defined. Further work is needed to determine whether SP from vaccination platforms induces similar but transient TYMP elevations, though current epidemiologic data do not link vaccines to increased lung cancer incidence. Future studies should use single-cell and spatial transcriptomic approaches to dissect how TYMP rewires epithelial, stromal, and immune interactions and to evaluate the therapeutic potential of TYMP inhibition in preventing fibrosis-induced oncogenesis.

In conclusion, this study reveals TYMP as a key molecular nexus through which SARS-CoV-2 SP promotes lung injury, fibrosis, and cancer development. By identifying TYMP as both a biomarker and effector of SP-induced pathology, our findings open new avenues for therapeutic intervention and highlight the urgent need for cancer risk assessment in individuals recovering from COVID-19.

## Acknowledgments

None.

## References

1. Zhu N, Zhang D, Wang W, Li X, Yang B, Song J, Zhao X, Huang B, Shi W, Lu R, Niu P, Zhan F, Ma X, Wang D, Xu W, Wu G, Gao GF, Tan W, China Novel Coronavirus I, Research T. A Novel Coronavirus from Patients with Pneumonia in China, 2019. N Engl J Med 2020; 382: 727–733.

2. John AE, Joseph C, Jenkins G, Tatler AL. COVID-19 and pulmonary fibrosis: A potential role for lung epithelial cells and fibroblasts. Immunol Rev 2021; 302: 228–240.

3. Zhao YM, Shang YM, Song WB, Li QQ, Xie H, Xu QF, Jia JL, Li LM, Mao HL, Zhou XM, Luo H, Gao YF, Xu AG. Follow-up study of the pulmonary function and related physiological characteristics of COVID-19 survivors three months after recovery. EClinicalMedicine 2020; 25: 100463.

4. Zhao Y, Yang C, An X, Xiong Y, Shang Y, He J, Qiu Y, Zhang N, Huang L, Jia J, Xu Q, Zhang L, Zhao J, Pei G, Luo H, Wang J, Li Q, Gao Y, Xu A. Follow-up study on COVID-19 survivors one year after discharge from hospital. Int J Infect Dis 2021; 112: 173–182.

5. Lee HY, Lee J, Lee CH, Han K, Choi SM. Risk of cancer incidence in patients with idiopathic pulmonary fibrosis: A nationwide cohort study. Respirology 2021; 26: 180–187.

6. Brown SW, Dobelle M, Padilla M, Agovino M, Wisnivesky JP, Hashim D, Boffetta P. Idiopathic Pulmonary Fibrosis and Lung Cancer. A Systematic Review and Meta-analysis. Ann Am Thorac Soc 2019; 16: 1041–1051.

7. Niu D, Song M, Chen M, Wu X, Zhang Y, Zhou R. COVID-19 vaccination status and the risk of developing lung diseases: A Mendelian randomization study. Medicine (Baltimore) 2025; 104: e43102.

8. Luo P, Liu J, Wang Z, Liao C, She L, Zou T, Chen J, Liu Z. Effects of COVID-19 vaccination on irAEs and prognosis in lung cancer patients receive PD-(L)1 inhibitors. Hum Vaccin Immunother 2025; 21: 2539593.

9. Chia SB, Johnson BJ, Hu J, Valenca-Pereira F, Chadeau-Hyam M, Guntoro F, Montgomery H, Boorgula MP, Sreekanth V, Goodspeed A, Davenport B, De Dominici M, Zaberezhnyy V, Schleicher WE, Gao D, Cadar AN, Petriz-Otano L, Papanicolaou M, Beheshti A, Baylin SB, Guarnieri JW, Wallace DC, Costello JC, Bartley JM, Morrison TE, Vermeulen R, Aguirre-Ghiso JA, Rincon M, DeGregori J. Respiratory viral infections awaken metastatic breast cancer cells in lungs. Nature 2025; 645: 496–506.

10. Roytenberg R, Yue H, DeHart A, Kim E, Bai F, Kim Y, Denning K, Kwei A, Zhang Q, Liu J, Zheng XL, Li W. Thymidine phosphorylase mediates SARS-CoV-2 spike protein enhanced thrombosis in K18-hACE2(TG) mice. Thrombosis research 2024; 244: 109195.

11. Rahman M, Irmler M, Keshavan S, Introna M, Beckers J, Palmberg L, Johanson G, Ganguly K, Upadhyay S. Differential Effect of SARS-CoV-2 Spike Glycoprotein 1 on Human Bronchial and Alveolar Lung Mucosa Models: Implications for Pathogenicity. Viruses 2021; 13.

12. Yonker LM, Swank Z, Bartsch YC, Burns MD, Kane A, Boribong BP, Davis JP, Loiselle M, Novak T, Senussi Y, Cheng CA, Burgess E, Edlow AG, Chou J, Dionne A, Balaguru D, Lahoud-Rahme M, Arditi M, Julg B, Randolph AG, Alter G, Fasano A, Walt DR. Circulating Spike Protein Detected in Post-COVID-19 mRNA Vaccine Myocarditis. Circulation 2023; 147: 867–876.

13. Ogata AF, Cheng CA, Desjardins M, Senussi Y, Sherman AC, Powell M, Novack L, Von S, Li X, Baden LR, Walt DR. Circulating Severe Acute Respiratory Syndrome Coronavirus 2 (SARS-CoV-2) Vaccine Antigen Detected in the Plasma of mRNA-1273 Vaccine Recipients. Clin Infect Dis 2022; 74: 715–718.

14. Li W, Yue H. Thymidine phosphorylase: A potential new target for treating cardiovascular disease. Trends Cardiovasc Med 2018; 28: 157–171.

15. Koukourakis MI, Giatromanolaki A, O’Byrne KJ, Comley M, Whitehouse RM, Talbot DC, Gatter KC, Harris AL. Platelet-derived endothelial cell growth factor expression correlates with tumour angiogenesis and prognosis in non-small-cell lung cancer. British journal of cancer 1997; 75: 477–481.

16. Li W, Gigante A, Perez-Perez MJ, Yue H, Hirano M, McIntyre TM, Silverstein RL. Thymidine phosphorylase participates in platelet signaling and promotes thrombosis. Circulation research 2014; 115: 997–1006.

17. Elamin YY, Rafee S, Osman N, KJ OB, Gately K. Thymidine Phosphorylase in Cancer; Enemy or Friend? Cancer Microenviron 2016; 9: 33–43.

18. Li W, Yue H. Thymidine Phosphorylase Is Increased in COVID-19 Patients in an Acuity-Dependent Manner. Front Med (Lausanne) 2021; 8: 653773.

19. Manne BK, Denorme F, Middleton EA, Portier I, Rowley JW, Stubben C, Petrey AC, Tolley ND, Guo L, Cody M, Weyrich AS, Yost CC, Rondina MT, Campbell RA. Platelet gene expression and function in patients with COVID-19. Blood 2020; 136: 1317–1329.

20. Russell CD, Valanciute A, Gachanja NN, Stephen J, Penrice-Randal R, Armstrong SD, Clohisey S, Wang B, Al Qsous W, Wallace WA, Oniscu GC, Stevens J, Harrison DJ, Dhaliwal K, Hiscox JA, Baillie JK, Akram AR, Dorward DA, Lucas CD. Tissue Proteomic Analysis Identifies Mechanisms and Stages of Immunopathology in Fatal COVID-19. Am J Respir Cell Mol Biol 2021.

21. Shimada M, Hasegawa H, Rikimaru T, Gion T, Hamatsu T, Yanashita Y, Shirabe K, Sugimachi K. The significance of thymidine phosphorylase activity in hepatocellular carcinoma and chronic diseased livers: a special reference to liver fibrosis and multicentric tumor occurrence. Cancer letters 2000; 148: 165–172.

22. Konda R, Sato H, Sakai K, Sato M, Orikasa S, Kimura N. Expression of platelet-derived endothelial cell growth factor and its potential role in up-regulation of angiogenesis in scarred kidneys secondary to urinary tract diseases. The American journal of pathology 1999; 155: 1587–1597.

23. Ikeguchi M, Sakatani T, Ueda T, Hirooka Y, Kaibara N. Thymidine phosphorylase activity in liver tissue and its correlation with multifocal occurrence of hepatocellular carcinomas. In vivo 2001; 15: 265–270.

24. Miller YE, Dwyer-Nield LD, Keith RL, Le M, Franklin WA, Malkinson AM. Induction of a high incidence of lung tumors in C57BL/6 mice with multiple ethyl carbamate injections. Cancer letters 2003; 198: 139–144.

25. Sozio F, Schioppa T, Sozzani S, Del Prete A. Urethane-induced lung carcinogenesis. Methods Cell Biol 2021; 163: 45–57.

26. Bongarzone S, Savickas V, Luzi F, Gee AD. Targeting the Receptor for Advanced Glycation Endproducts (RAGE): A Medicinal Chemistry Perspective. Journal of medicinal chemistry 2017; 60: 7213–7232.

27. Grommes J, Soehnlein O. Contribution of neutrophils to acute lung injury. Mol Med 2011; 17: 293–307.

28. Portales AE, Mustafá ER, McCarthy CI, Cornejo MP, Couto PM, Gironacci MM, Caramelo JJ, Perelló M, Raingo J. ACE2 internalization induced by a SARS-CoV-2 recombinant protein is modulated by angiotensin II type 1 and bradykinin 2 receptors. Life Sci 2022; 293: 120284.

29. Lu Y, Zhu Q, Fox DM, Gao C, Stanley SA, Luo K. SARS-CoV-2 down-regulates ACE2 through lysosomal degradation. Mol Biol Cell 2022; 33: ar147.

30. Bartolomé A, Liang J, Wang P, Ho DD, Pajvani UB. Angiotensin converting enzyme 2 is a novel target of the γ-secretase complex. Scientific Reports 2021; 11: 9803.

31. Affandi KA, Tizen NMS, Mustangin M, Zin R. p40 Immunohistochemistry Is an Excellent Marker in Primary Lung Squamous Cell Carcinoma. J Pathol Transl Med 2018; 52: 283–289.

32. Bishop JA, Teruya-Feldstein J, Westra WH, Pelosi G, Travis WD, Rekhtman N. p40 (ΔNp63) is superior to p63 for the diagnosis of pulmonary squamous cell carcinoma. Modern Pathology 2012; 25: 405–415.

33. Gurley KE, Moser RD, Kemp CJ. Induction of Lung Tumors in Mice with Urethane. Cold Spring Harb Protoc 2015; 2015: pdb.prot077446.

34. Handa M, Li W, Morioka K, Takamori A, Yamada N, Ihaya A. Adventitial delivery of platelet-derived endothelial cell growth factor gene prevented intimal hyperplasia of vein graft. J Vasc Surg 2008; 48: 1566–1574.

35. Li W, Tanaka K, Morioka K, Uesaka T, Yamada N, Takamori A, Handa M, Tanabe S, Ihaya A. Thymidine phosphorylase gene transfer inhibits vascular smooth muscle cell proliferation by upregulating heme oxygenase-1 and p27KIP1. Arteriosclerosis, thrombosis, and vascular biology 2005; 25: 1370–1375.

36. Du LL YH, Rorabaugh BR, Li O, DeHart AR, Toloza-Alvarez G, Hong L, Denvir J, Thompson E, and Li W. Thymidine phosphorylase deficiency or inhibition preserves cardiac function in mice with acute myocardial infarction. Journal of the American Heart Association (In press) 2023.

37. Yue H, Tanaka K, Furukawa T, Karnik SS, Li W. Thymidine phosphorylase inhibits vascular smooth muscle cell proliferation via upregulation of STAT3. Biochimica et biophysica acta 2012; 1823: 1316–1323.

38. Turnbull IR, Fuchs A, Remy KE, Kelly MP, Frazier EP, Ghosh S, Chang SW, Mazer MB, Hess A, Leonard JM, Hoofnagle MH, Colonna M, Hotchkiss RS. Dysregulation of the leukocyte signaling landscape during acute COVID-19. PloS one 2022; 17: e0264979.

39. Chakraborty D, Šumová B, Mallano T, Chen C-W, Distler A, Bergmann C, Ludolph I, Horch RE, Gelse K, Ramming A, Distler O, Schett G, Šenolt L, Distler JHW. Activation of STAT3 integrates common profibrotic pathways to promote fibroblast activation and tissue fibrosis. Nature Communications 2017; 8: 1130.

40. Hong L, Yue H, Cai D, DeHart A, Toloza-Alvarez G, Du L, Zhou X, Fan X, Huang H, Chen S, Rahaman SO, Zhuang J, Li W. Thymidine Phosphorylase Promotes Abdominal Aortic Aneurysm via VSMC Modulation and Matrix Remodeling in Mice and Humans. Cardiovasc Ther 2024; 2024: 1129181.

41. Ogarek N, Oboza P, Olszanecka-Glinianowicz M, Kocelak P. SARS-CoV-2 infection as a potential risk factor for the development of cancer. Front Mol Biosci 2023; 10: 1260776.

42. Jahankhani K, Ahangari F, Adcock IM, Mortaz E. Possible cancer-causing capacity of COVID-19: Is SARS-CoV-2 an oncogenic agent? Biochimie 2023; 213: 130–138.

43. Tang X, Lu L, Li X, Huang P. Bridging Cancer and COVID-19: The Complex Interplay of ACE2 and TMPRSS2. Cancer Med 2025; 14: e70829.

44. Kunvariya AD, Dave SA, Modi ZJ, Patel PK, Sagar SR. Exploration of multifaceted molecular mechanism of angiotensin-converting enzyme 2 (ACE2) in pathogenesis of various diseases. Heliyon 2023; 9: e15644.

45. Pedroza M, Le TT, Lewis K, Karmouty-Quintana H, To S, George AT, Blackburn MR, Tweardy DJ, Agarwal SK. STAT-3 contributes to pulmonary fibrosis through epithelial injury and fibroblast-myofibroblast differentiation. Faseb j 2016; 30: 129–140.

46. Yue H, Li W, Desnoyer R, Karnik SS. Role of nuclear unphosphorylated STAT3 in angiotensin II type 1 receptor-induced cardiac hypertrophy. Cardiovasc Res 2010; 85: 90–99.

47. Wen Z, Liu T, Xu X, Acharya N, Shen Z, Lu Y, Xu J, Guo K, Shen S, Zhao Y, Wang P, Li S, Chen W, Li H, Ding Y, Shang M, Guo H, Hou Y, Cui B, Shen M, Huang Y, Pan T, Qingqing W, Cao Q, Wang K, Xiao P. Interleukin-16 enhances anti-tumor immune responses by establishing a Th1 cell-macrophage crosstalk through reprogramming glutamine metabolism in mice. Nature Communications 2025; 16: 2362.

48. Ke B, Wei T, Huang Y, Gong Y, Wu G, Liu J, Chen X, Shi L. Interleukin-7 Resensitizes Non-Small-Cell Lung Cancer to Cisplatin via Inhibition of ABCG2. Mediators Inflamm 2019; 2019: 7241418.

49. Malik A, Kanneganti TD. Function and regulation of IL-1α in inflammatory diseases and cancer. Immunol Rev 2018; 281: 124–137.

50. Belcher A, Zulfiker AHM, Li OQ, Yue H, Gupta AS, Li W. Targeting Thymidine Phosphorylase With Tipiracil Hydrochloride Attenuates Thrombosis Without Increasing Risk of Bleeding in Mice. Arteriosclerosis, thrombosis, and vascular biology 2021; 41: 668–682.

