## Supplemental materials for "Thymidine Phosphorylase Drives SARS-CoV-2 Spike Protein-Induced Lung Tumorigenesis"

**Supplemental Table 1. Antibodies used in the project**

| Antibodies used | Applications | Concentration used |
|---|---|---|
| Anti-RAGE antibody, ab30381 | IHC | 1:50 |
| Anti-myeloperoxidase antibody [EPR20257] ab208670 | IHC | 1:100 |
| Myc-Tag (9B11) mouse monoclonal antibody #2276 | WB | 1:1000-3000 |
| ACE2 antibody #4355 | WB | 1:1000 |
| GFP (D5.1) rabbit monoclonal antibody #2956 | WB | 1:1000 |
| Pan-Actin (D18C11) rabbit monoclonal antibody (HRP Conjugate) #12748 | WB | 1:3000-5000 |
| C9 tag polyclonal antibody, PAB26959 | WB | 1:500-1000 |
| Anti-p40 - DeltaNp63 antibody [EPR17863-47], ab203826 | IHC | 1:100 |
| Phospho-Stat3 (Tyr705) (D3A7) rabbit monoclonal antibody #9145 | IHC | 1:100 |
| Stat3, BD610190, mouse monoclonal antibody | IHC | 1:100 |
| CD68 antibody \| FA-11, MCA1957 | IHC | 1:100 |
| Rb @ PECAM-1 (CD31), 11265-1-AP | IHC | 1:200 |
| Anti-α-smooth muscle actin (ACTA2) antibody, A2547 | IHC | 1:200 |
| Anti-rabbit IgG, HRP-linked antibody #7074 | WB | 1:2000-5000 |
| Anti-mouse IgG, HRP-linked antibody #7076 | WB | 1:2000-5000 |

| | | |
|---|---|---|
| ImmPRESS-AP Horse Anti-Rabbit IgG Polymer Detection Kit, Alkaline Phosphatase, | IHC | N/A |
| ImmPRESS-AP Horse Anti-Mouse IgG Polymer Detection Kit, Alkaline Phosphatase, | IHC | N/A |
| M.O.M. (Mouse on Mouse) ImmPRESS HRP (Peroxidase) Polymer Kit | IHC | N/A |
| Vector Red Substrate Kit, Alkaline Phosphatase (AP) | IHC | N/A |
| Vector Blue Substrate Kit, Alkaline Phosphatase (AP) | IHC | N/A |

**Supplemental Table 2. Matched cohort sizes and baseline demographics (after PSM)**

| Comparison | Cohort A (definition) | N analyzed | Cohort B (definition) | N analyzed | Age, mean (SD) | Female, % | White, % | Not Hispanic/Latino, % |
|---|---|---|---|---|---|---|---|---|
| Current smokers: COVID-19 vs No COVID-19 | COVID-19, unvaccinated | 171,671 | No COVID-19, vaccinated | 171,671 | 54.1 (15.0) | 46.1 | 65.5 | 84.3 |
| Former smokers: COVID-19 vs No COVID-19 | COVID-19, unvaccinated | 425,056 | No COVID-19, vaccinated | 425,056 | 62.4 (15.1) | 46.5 | 76.3 | 85.3 |
| Never smokers: COVID-19 vs No COVID-19 | COVID-19, vaccinated | 1,704,065 | No COVID-19, vaccinated | 1,710,920 | 45.8 (21.5) | 56.5 | 55.8 | 73.3 |
| Current smokers with COVID-19: Unvaccinated vs Vaccinated | Unvaccinated | 199,981 | Vaccinated | 199,981 | 56.4 (15.4) | 51.1 | 66.1 | 85.9 |
| Former smokers with COVID-19: Unvaccinated vs Vaccinated | Unvaccinated | 519,183 | Vaccinated | 519,183 | 64.6 (15.8) | 50.1 | 77.0 | 86.9 |

**Supplemental Table 3. Whole blood cell counts in mice treated with SARS-CoV-2 Spike protein-containing cell lysate.**

| | **hACE2$^{TG}$** | **hACE2$^{TG}$/*Tymp*$^{-/-}$** | P value |
|---|---|---|---|
| WBC (K/uL) | 2.987 ± 0.707 | 1.867 ± 0.094 | 0.191311 |
| NE (K/uL) | 1.110 ± 0.175 | 1.370 ± 0.349 | 0.542268 |
| LY (K/uL) | 1.320 ± 0.655 | 0.7000 ± 0.07 | 0.399928 |
| MO (K/uL) | 0.5033 ± 0.091 | 0.1300 ± 0.032 | 0.017797 |
| EO (K/uL) | 0.03667 ± 0.027 | 0.0033 ± 0.0033 | 0.291974 |
| BA (K/uL) | 0.01000 ± 0.010 | 0.000 ± 0.000 | 0.373901 |
| NE % | 39.66 ± 8.384 | 55.40 ± 1.201 | 0.136585 |
| LY % | 40.03 ± 13.892 | 37.36 ± 2.850 | 0.859825 |
| MO % | 19.05 ± 5.867 | 6.937 ± 1.694 | 0.118291 |
| EO % | 1.000 ± 0.59 | 0.2733 ± 0.015 | 0.285745 |
| BA % | 0.2600 ± 0.166 | 0.023 ± 0.023 | 0.231837 |
| RBC (M/uL) | 7.917 ± 0.137 | 8.040 ± 0.053 | 0.448703 |
| Hb (g/dL) | 10.13 ± 0.088 | 11.13 ± 0.24 | 0.017458 |
| HCT % | 29.30 ± 0.3 | 33.37 ± 0.41 | 0.001318 |
| MCV (fL) | 37.03 ± 0.273 | 41.53 ± 0.233 | 0.000233 |
| MCH (pg) | 12.80 ± 0.100 | 13.87 ± 0.318 | 0.032901 |
| MCHC (g/dL) | 34.57 ± 0.12 | 33.37 ± 0.825 | 0.223591 |
| RDW % | 17.10 ± 0.231 | 17.00 ± 0.346 | 0.821989 |
| PLT (K/uL) | 605.0 ± 16.773 | 517.0 ± 11.358 | 0.012212 |

| | | | |
|---|---|---|---|
| MPV (fL) | 4.500 ± 0.3 | 3.967 ± 0.033 | 0.151987 |

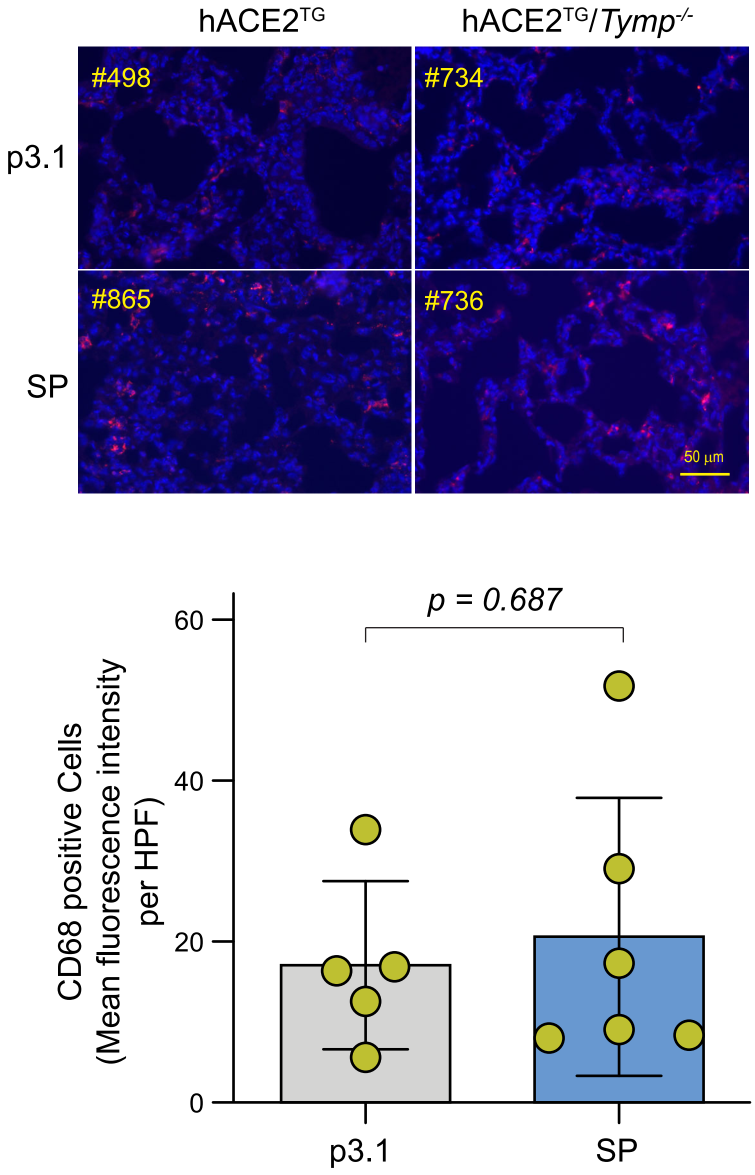


**Supplemental Figure 1.** K18-hACE2TG (hACE2TG) and K18-hACE2TG/*Tymp*$^{-/-}$ (hACE2TG/*Tymp*$^{-/-}$) mice were treated with P3.1 or SP via intratracheal administration. Lungs were harvested 24 hours later, and lung sections were stained for CD68, a macrophage marker. Images were visualized using Alexa Fluor 568. Mean fluorescence intensity of Alexa fluor 568 was quantified using ImageJ as a measure of macrophage infiltration.

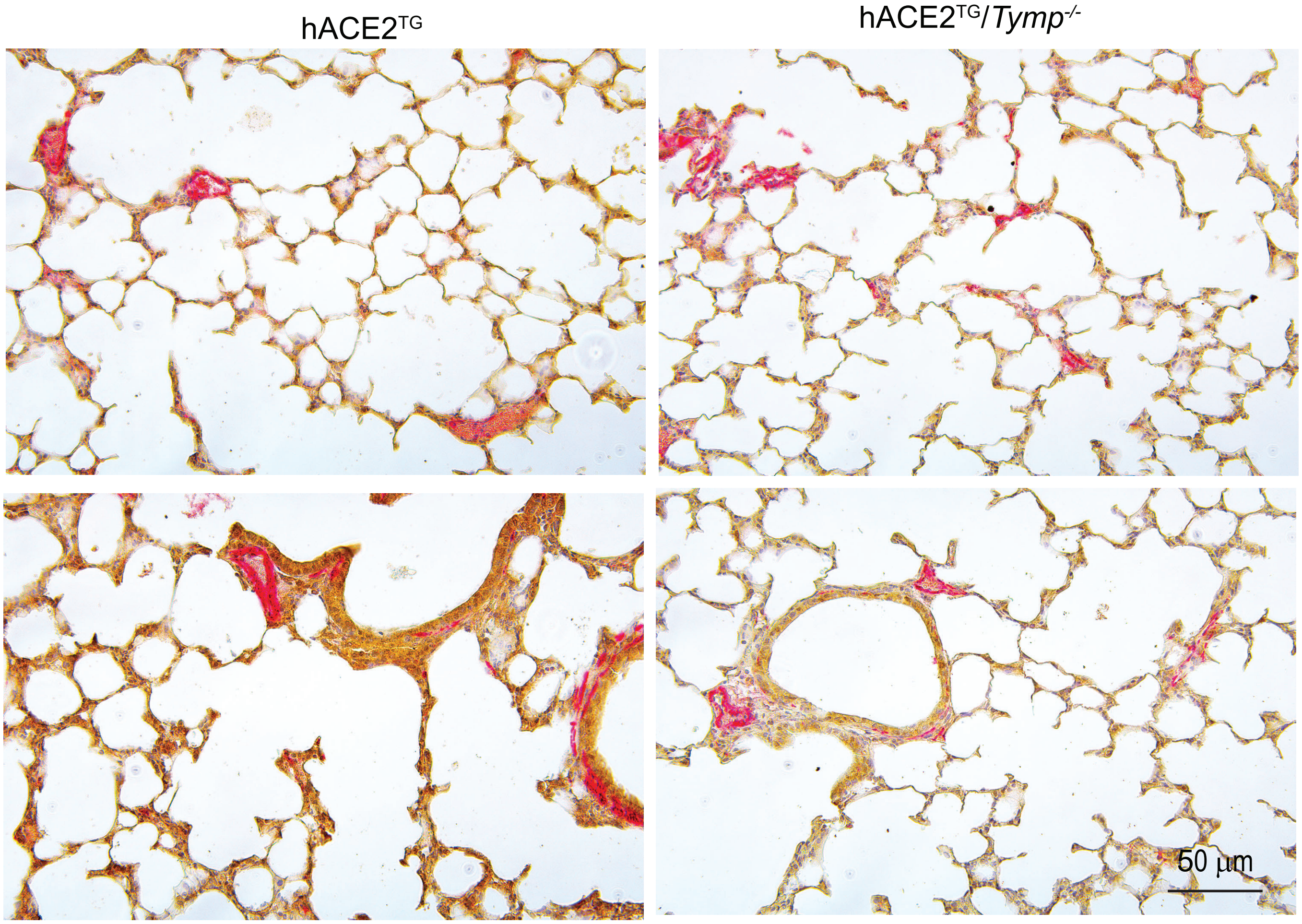


**Supplemental Figure 2.** Lung sections from K18-hACE2TG (hACE2TG) and K18-hACE2TG/*Tymp*$^{-/-}$ (hACE2TG/*Tymp*$^{-/-}$) mice were double stained for CD31, a marker of endothelial cells, and alpha-smooth muscle actin (α-SMA), a marker of vascular smooth muscle cells. Nuclei were counterstained with hematoxylin.

**Supplemental Figure 3. Cytokine Array Assay**

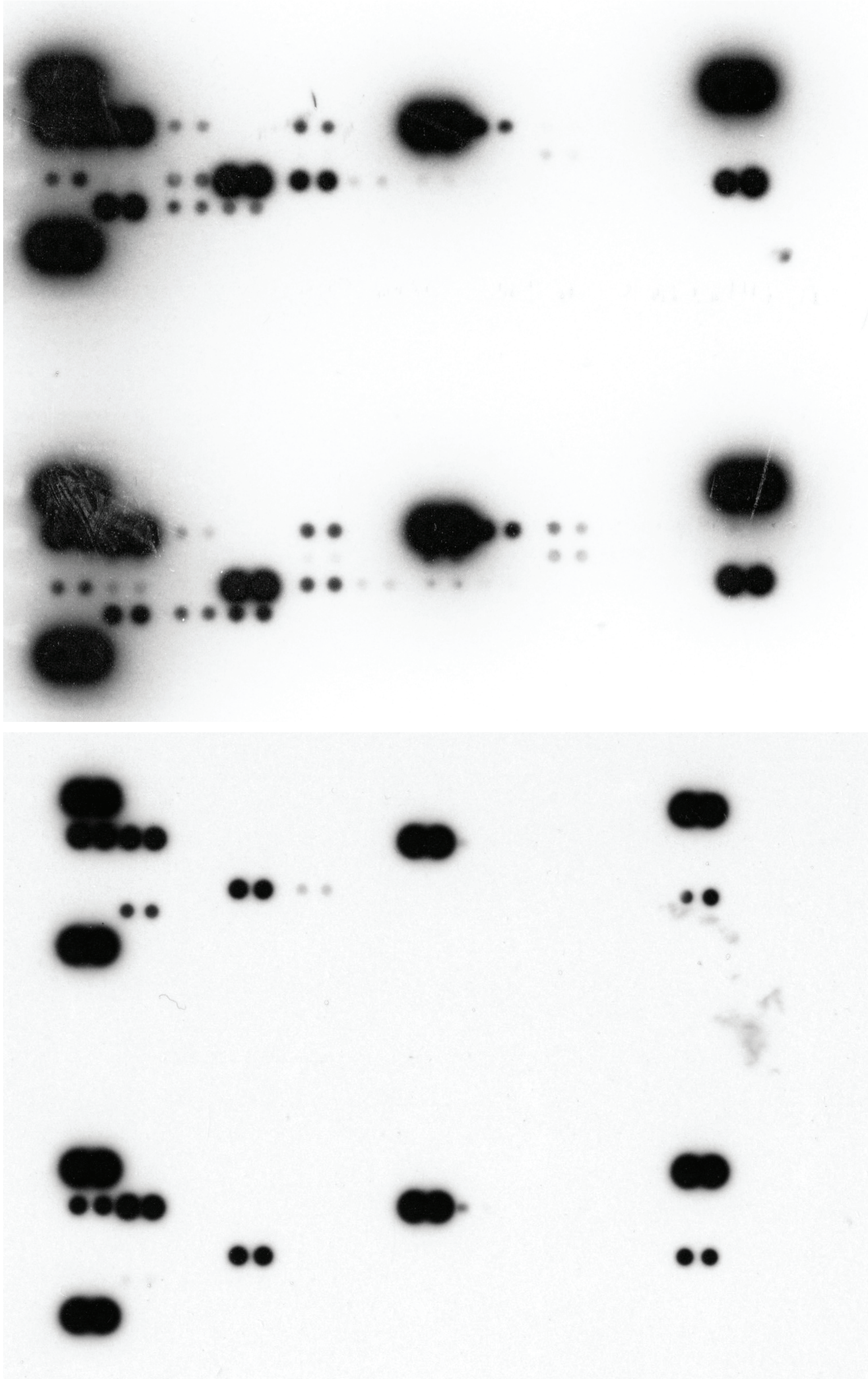

#281
K18-hACE2$^{TG}$

#282
K18-hACE2$^{TG}$/*Tymp*$^{-/-}$

*ECL*
*Pico*
*20 seconds*

#281
K18-hACE2$^{TG}$

#282
K18-hACE2$^{TG}$/*Tymp*$^{-/-}$

*ECL*
*Pico*
*5 seconds*
